# A targeted and tuneable DNA damage tool using CRISPR/Cas9

**DOI:** 10.1101/2021.01.23.427907

**Authors:** Ioannis Emmanouilidis, Natalia Fili, Alexander W. Cook, Yukti Hari-Gupta, Ália dos Santos, Lin Wang, Marisa Martin-Fernandez, Peter J.I. Ellis, Christopher P. Toseland

**Author notes:** Current address: MRC LMCB, University College London, London, WC1E 6BT, UK.

## Abstract

Mammalian cells are constantly subjected to a variety of DNA damaging events that lead to the activation of DNA repair pathways. Understanding the molecular mechanisms of the DNA damage response allows the development of therapeutics which target elements of these pathways.

Double-Strand Breaks (DSB) are particularly deleterious to cell viability and genome stability. Typically, DSB repair is studied using DNA damaging agents such as ionising irradiation or genotoxic drugs. These induce random lesions at non-predictive genome sites, where damage dosage is difficult to control. Such interventions are unsuitable for studying how different DNA damage recognition and repair pathways are invoked at specific DSB sites in relation to the local chromatin state.

The RNA-guided Cas9 (CRISPR associated protein 9) endonuclease enzyme, is a powerful tool to mediate targeted genome alterations. Cas9-based genomic intervention is attained through DSB formation in the genomic area of interest. Here, we have harnessed the power to induce DSBs at defined quantities and locations across the human genome, using custom-designed promiscuous guide RNAs, based on *in silico* predictions. This was achieved using electroporation of recombinant Cas9-guide complex which provides a generic, low-cost and rapid methodology for inducing controlled DNA damage in cell culture models.

## INTRODUCTION

One of the most critical processes for living organisms is to maintain genome integrity using DNA damage surveillance and repair mechanisms. These mechanisms prevent cells from progressing through cell division, which would propagate the defective genome to daughter cells ^1^. If lesions are not repaired, mutations can accumulate leading to cell senescence, ageing and the onset of disease such as cancer.

Approximately 10-50 double strand breaks (DSBs) occur per cell cycle in human cells ^2,3^. DSBs are considered one of the most genotoxic types of DNA damage because both DNA strands are severed, and thus any error in correctly re-joining the broken ends may lead to insertions, translocations, deletions and chromosome fusions that further promote genome instability^4^. Multiple competing repair programmes exist in order to repair such lesions, including error-free (homologous recombination, HR) and error prone (non-homologous end joining, NHEJ; mismatch-mediated end joining, MMEJ) pathways ^4^. The choice of which repair pathway is invoked depends on the nature of the break, on the local chromatin context, and on cell cycle stage. When the nature or level of DNA damage is beyond repair, apoptotic mechanisms are activated ^5^. This apoptotic response is frequently exploited by cytotoxic drugs such as bleomycin or cisplatin, which induce severe aberrations within the DNA structure. Understanding the mechanisms involved in pathway choice for DSB repair is crucial to develop targeted therapeutic interventions in cancer cells, as for example is the case of PARP inhibitors in Brca1-deficient cancers ^6^.

In order to study DSB response mechanisms, damage must first be induced. To date, this has largely been achieved using untargeted genotoxic drugs and/or irradiation. While this allows some definition of how DSB repair varies according to cell type, it does not allow interrogation of how repair is influenced by local chromatin states, which can alter upon damage ^7^. More targeted approaches have also been developed for the study of DSBs, reliant upon the inducible expression of restriction endonucleases that sever the DNA helix at specific locations. These experimental systems encompass the use of *Ppol*^8^, *SceI*^9^ and *AsiSI* enzymes^10^ – each of which cuts the DNA in different locations, and which may be introduced to cells either transiently, or via genomic incorporation of an inducible transgene. In particular, inducible expression of *AsiSI* using the DiVA has cell line been instrumental in beginning to unpick the complexity of DNA repair pathway choice in different genomic contexts ^10^.

However, each of these systems only addresses a very limited selection of chromosome regions due to the requirement for specific restriction enzyme cut sites. *SceI* is a meganuclease with no endogenous recognition site in mammalian genomes, and thus its target sequence must be transgenically introduced into the desired location in the genome. *PpoI* has several recognition sites within ribosomal RNA repeats and thus only allows the study of the nucleolar DNA damage response. *AsiSI* is methylation-sensitive and cuts at non-methylated CpG-rich sequences, which are primarily located at promoters and enhancers^10^. There is an urgent need within the field to expand the toolbox to allow wider interrogation of DNA responses in different chromatin contexts, and – equally importantly – different cell types.

Therefore, we wished to design a methodology to induce a tuneable number of DSBs in a broad range of genomic contexts. This can be used to probe the context specificity of DNA damage responses and pathway choice. Programmable endonucleases such as Transcription activator-like effector nucleases (TALENS) or Zinc finger nucleases (ZFNs), and CRISPR-Cas9 have all been widely used to allow DSB induction at arbitrary genomic loci. However, typically these programmed approaches are designed to generate a single DSB at a single specific site for the purpose of gene targeting, with broader induction of DNA damage seen as a disadvantage^11–15^. Here, in contrast, we use electroporation of recombinant Cas9 and synthetic promiscuous guide RNAs to introduce multiple DSBs in mammalian cell culture, with both the number of breaks and the desired target context being tuneable (Figure 1A). This will allow us to assess the DNA damage response across a wide range of damage severity, at specified genomic locations, in any electroporatable cell type.

**Figure 1.**
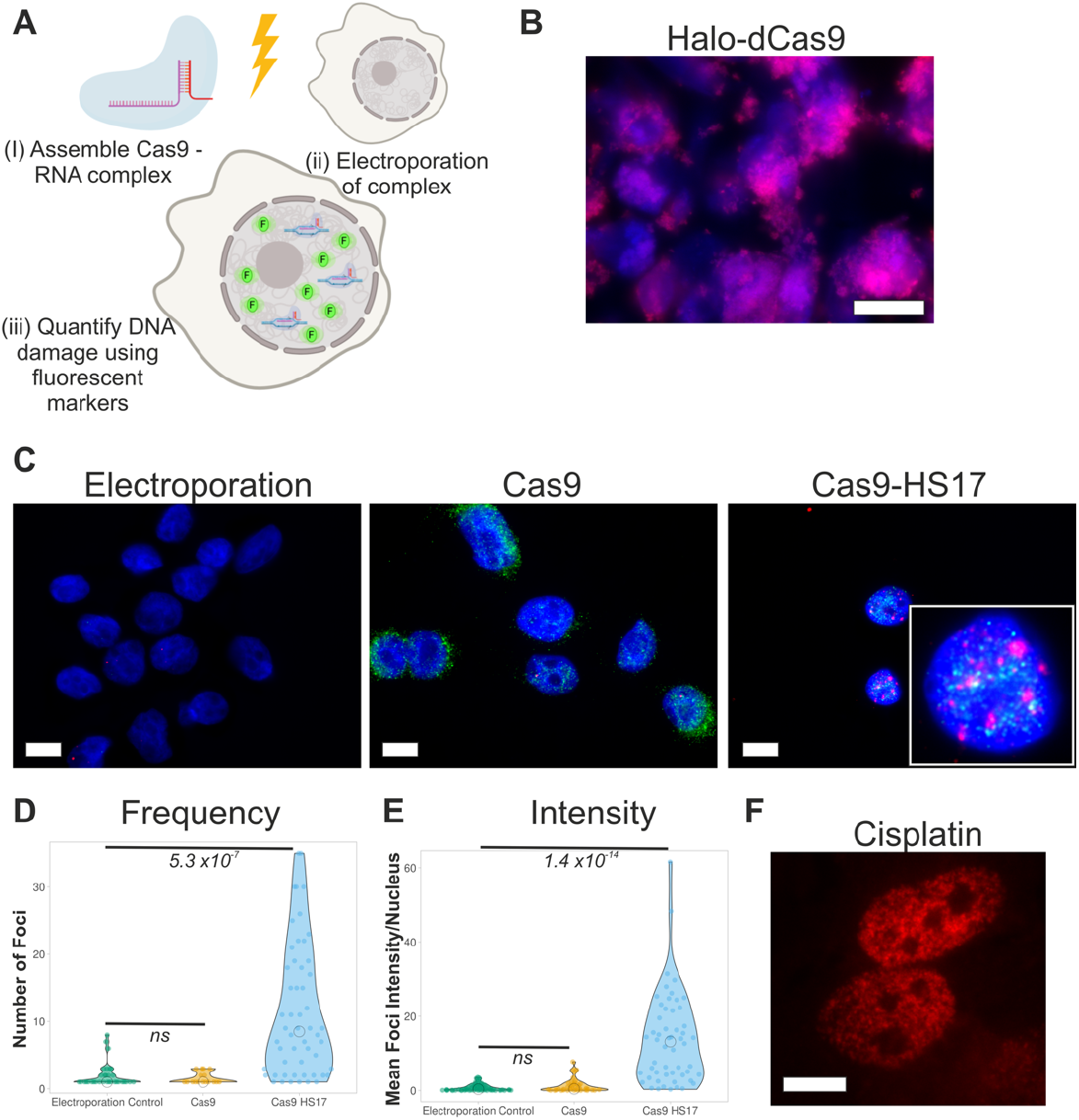
Electroporation of Cas9 to induce DNA damage. (A) Cartoon depicting the methodology. Recombinant Cas9 is bound to the crRNA and tracrRNA (i). The complex is then electroporated into the mammalian cell line (ii). DNA damage is then detected and quantified using fluorescent markers (iii). (B) Example widefield image of Halo-dCas9 in MCF10a cells stained with TMR ligand (magenta) and Hoechst for DNA (blue). Scale bar is 10 μm. (C) Immunofluorescence staining against Cas9 (green) and γH2AX (red) in MCF10a cells. Electroporation is a control for background signals. Cas9 refers to electroporation of only Cas9. HS17 is the guide RNA predicted to cut the genome at 17 locations. DNA was stained with Hoechst (blue). Scale bar is 10 μm. (D-E) Quantitative image analysis from the experiments in (C). The Number of Foci per nucleus and Mean Foci Intensity per nucleus are plotted for each condition. Each data point is an individual nucleus. The p value is calculated from a two-tailed t-test. (F) Immunofluorescence staining of γH2AX in MCF10a cells following treatment with 25 μM cisplatin for 4 hrs.

## RESULTS

### Proof of concept using electroporation of recombinant Cas9

Our approach utilises electroporation of recombinant Cas9 instead of traditional transfection or engineering stable expression within cell line models. This reduces the perturbation to the cell line, removes the time delay to express Cas9 and enables multiple cells lines to be readily studied. We first established the electroporation efficiency with nuclease-deficient Cas9 (dCas) to avoid any complications arising from non-specific nuclease activity. We electroporated Halo-dCas9 in to MCF10a cells and then stained the cells with TMR halo-ligand. On average, we observed 40 % of cells contained dCas9 (Figure 1B).

We then assessed the ability of wild type Cas9-RNA complex to be electroporated and induce DNA damage. For this, we used a previously designed promiscuous crRNA guide to induced up to 17 DSBs (HS17) ^16^. The Cas9-RNA complex was pre-formed and then electroporated into MCF10a cells. Immunofluorescence staining against Cas9 marked electroporated cells and γH2AX staining was used as a read-out of DNA damage. The latter was quantified in terms of number of foci and total nuclear intensity. Electroporation without guide RNA did not induce DNA damage. However, DNA damage was induced through electroporation of the Cas9-RNA complex (Figure 1C-E), as occurred following exposure of the cells to cisplatin (Figure 1F). Overall, we conclude that this approach is viable for inducing DNA damage, independent of cell line engineering.

### Design of promiscuous guide RNAs

Having shown that the electroporation method is feasible, we set out to design a series of promiscuous guide RNAs to induce DNA damage across a range of sites. The FlashFry software ^17^ compresses the genome into an organised index for easier and faster discovery of target sequences. This was used to identify 23bp (crRNA/PAM + sequence) target sites in the human genome. A total of five sequences were selected from the generated output list (Table 1). Sequences were selected based on the prediction that each would cut at 50, 100 and 150 locations within the human genome, with a PAM sequence matching *n*GG or *n*GA. There is consensus between the predicted cuts using FlashFry and Ensembl (Table 1). Specifically, there are two versions for each of the 50 and 100 crRNA sequences. These target relatively GC-rich sequences 50A (60.9% GC) and 100A (56.5% GC), or AT-rich regions 50B (69.6% AT) and 100B (60.9% AT). A quantified overview of the hits per chromosome is summarised in Supplementary Figure 1, including hits relative to the number of bases for a given chromosome. The hits are randomly distributed across the chromosomes but not all chromosomes are targeted with each guide. In addition to these, we used the previously-designed HS17 gRNA (see above).

**Table 1.**
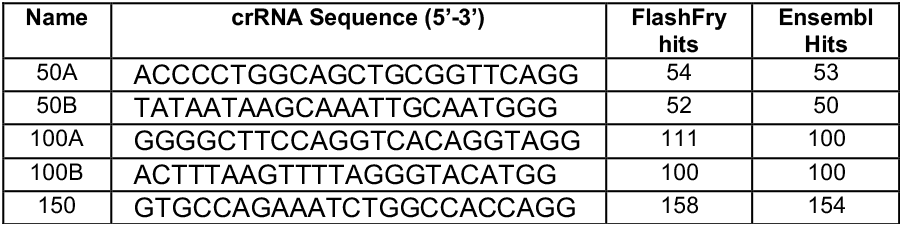
Candidate sequences for promiscuous targeting of the human genome. Predicted number of hits are presented from FlashFry and Ensembl.

### Tuning the number of DNA damage sites using Cas9 and promiscuous guides

#### (I) Single particle analysis of DNA damage foci

As in the proof-of-concept experiment, the formation of γH2AX foci was used as a marker of DNA damage to report upon the activity of Cas9 in complex with the guide RNAs following electroporation. Electroporation alone, or with only Cas9, showed minimal γH2AX staining and foci formation (Figure 2).

**Figure 2.**
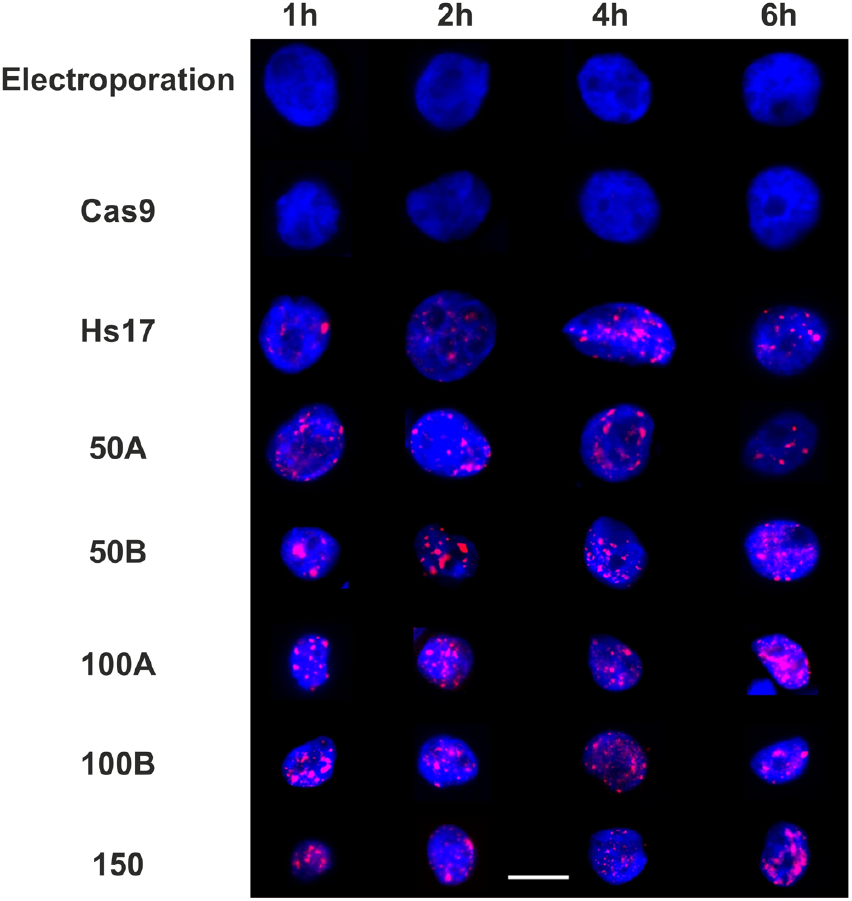
Time course of Cas9-induced DNA damage. Example widefield images of MCF10a cells stained for DNA with Hoechst (blue) and γH2AX (red). Electroporation is a control for background signals. Cas9 refers to electroporation of Cas9 alone. HS17 is the crRNA predicted to cut the genome at 17 locations16, while 50A/B, 100A/B and 150 are our designed promiscuous crRNA which cut at 50, 100 and 150 predicted sites, respectively. ‘A’ versions are GC-selective sequences while ‘B’ versions are AT-selective. The timing is measured from electroporation onwards. Scale bar is 10 μm.

Upon electroporation of the Cas9:RNA complex, formation of γH2AX foci can be observed from one-hour following electroporation (Figure 2) through the six-hour observation window. These results confirm that the promiscuous guides are viable for the induction of DSBs through Cas9 nuclease activity.

To investigate the induction of DSBs quantitatively, we used single particle image analysis on wide-field images of nuclei. Consistent with Figure 2, the majority of the cells in the control measurements revealed minimal DNA damage for all time courses (mean foci cell^−1^ = 1.3). Electroporation of the Cas9:RNA complexes led to a significant accumulation of γH2AX foci for all guide RNA, except centromere, compared to controls (Figure 1D, 2).

Despite the presence of foci 1-hour post-electroporation, the number of foci detected varies across the time course between the different guide RNAs. For example, the mean number of foci with 100B electroporated cells remains constant at approximately 15.5 foci cell^−1^ throughout the time course, while in 100A electroporated cells, the number of foci increased up to 2 hours and then decreased in the following 4 hours. However, all of the conditions are capable of retaining a large number of γH2AX foci within the 6-hour time frame because either DNA damage repair is incomplete or Cas9 remains active. The temporal difference is potentially due to variations in cut and repair efficiencies within the different genomic regions.

Using guide RNAs with increasing predictive cuts lead to a broader distribution in the number of γH2AX foci (Figure 3). However, the overall cell response to the cuts shows that only a small population of electroporated cells are capable of reaching the predicted number of cuts, such as those electroporated with HS17 (30%) and few (7%) in the case of 50A and 50B (Figure 3). This may relate to the accessibility of the sites to nuclease activity. Occasionally, there are cells like in HS17, 50A and 50B that display twice the number of predicted cuts, suggesting that these are cells in G2/M phase with duplicated genomes. The are no trends across the time courses between AT or GC sequence bias. For example, there are differences between 50A and 50B at 4 hours, AT-selective 50B guide generates more foci. Whereas, the GC-selective 100A has more foci at 2 hours.

**Fig.3.**
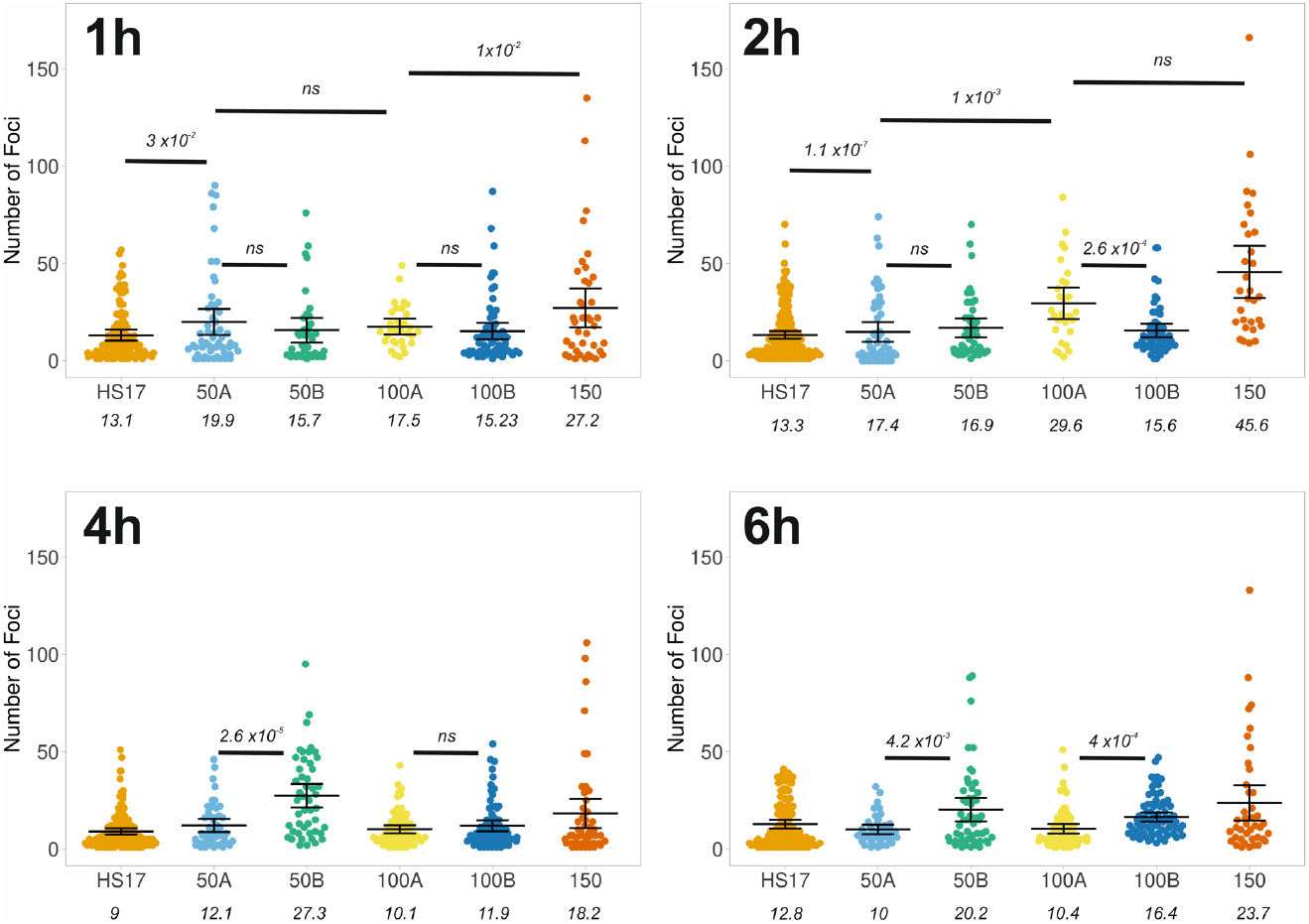
Quantification of DNA damage foci. Quantitative image analysis, as described in the methods, from the experiments in Figure 2. Foci were determined based on staining of γH2AX. The Number of Foci per nucleus is plotted for each crRNA (Table 1). Each data point is an individual nucleus. The p value is calculated from a two-tailed t-test and the error bars represent standard deviation. The mean values are presented.

Caution must be exercised when attempting to count absolute values when a high density of foci needs to be measured in a spatially and genomically confined region, and through the use of widefield imaging in a single focal plane.

To address these technical limitations and to provide further validation of our approach, we performed confocal microscopy to extract a 3D volume to better quantify the number of foci. This enabled us to correctly quantify particles that overlap in 2C but are separated in 3D, although the overall spatial resolution remains similar to wide field imaging. We used the 50A and 50B guides at the 2-hour measurements (Figure 4A-D). This time point was chosen because it was possible to resolve statistically significant differences between the predicted 17, 50 and 100 cuts. The γH2AX foci were well resolved and clearly visible throughout the nuclear body (Figure 4B and 4D). Particle analysis was then performed across the 3D stack and, as expected, we resolved more foci for each guide (Figure 4E). Specifically, we observed a mean of 49 and 35 foci cell^−1^ for 50A and 50B, respectively, compared to 17 and 16 foci cell^−1^ observed in conventional wide-field microscopy. Thus, when resolved by confocal stacking, the average foci count is significantly closer to the expected number of cuts. In the confocal analysis, our ability to detect a high number of particles is not a limiting factor, as shown by applying the same confocal approach with cells treated with cisplatin (Figure 4F) where we detect an average of 136 foci.

**Fig.4.**
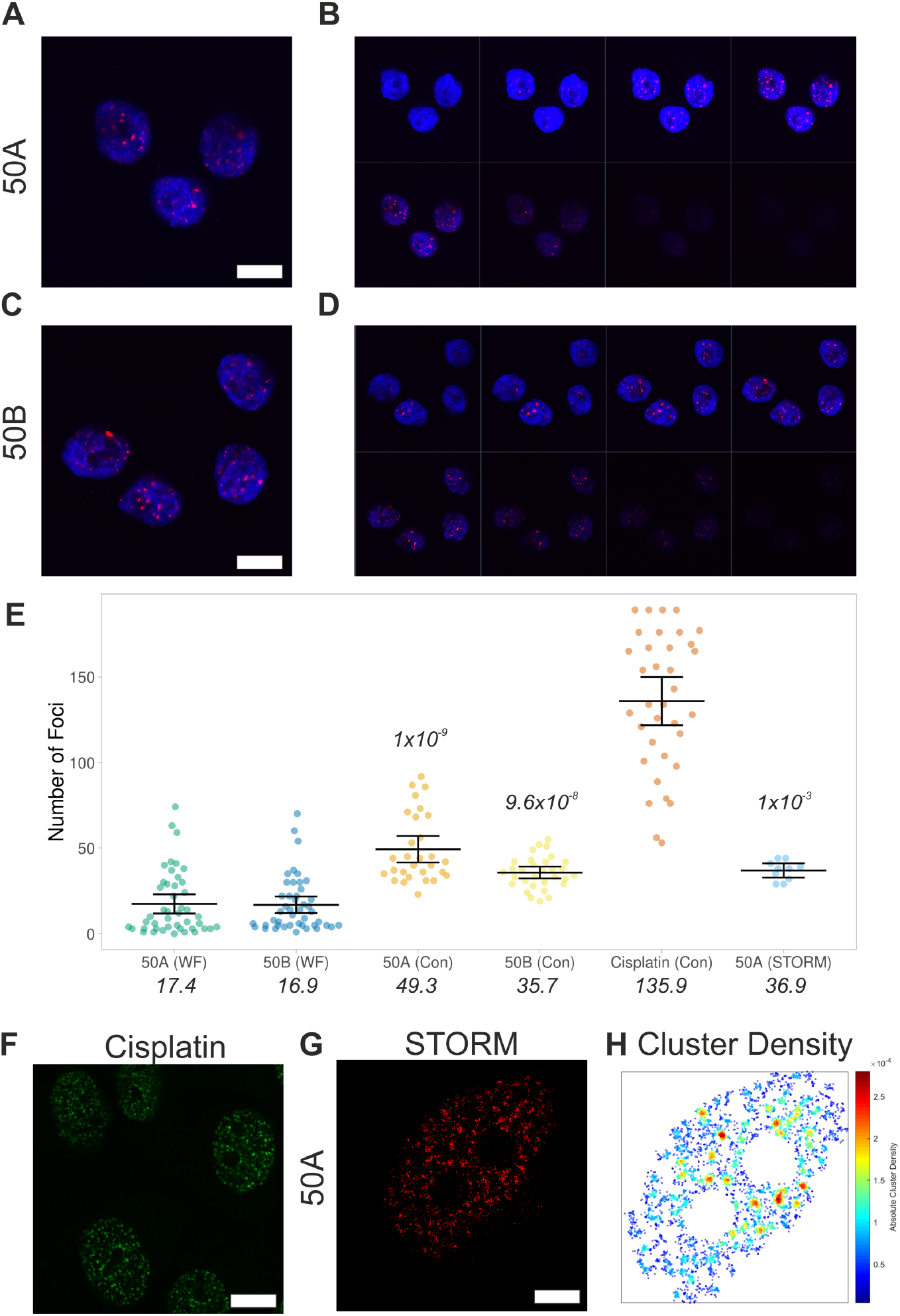
Impact of imaging methods upon the quantification of γH2AX foci. (A-B) Confocal image and nuclear z-stack at 500 nm intervals of MCF10a cells stained for DNA with Hoechst (blue) and γH2AX (red), 2 hrs after electroporation with Cas9 and 50A guide (GC-selective). Scale bar: 10 μm. (C-D) The equivalent measurement with AT-selective 50B guide. Scale bar: 10 μm. (E) Quantitative image analysis to determine the Number of Foci per nucleus. ‘WF’ refers to standard widefield imaging, as shown in Figure 2. ‘Con’ denotes confocal imaging, as shown in A-D. ‘STORM’ refers to the number of foci determined by super resolution imaging and cluster analysis in panels F-G. A comparison is shown for MCF10a cells treated with 25 μM cisplatin for 4 hrs. Each data point is an individual nucleus. The p value is calculated from a two-tailed t-test comparing measurements to the standard widefield imaging and the error bars represent sd. The mean values are presented. (F) Confocal image of γH2AX in MCF10a cells following treatment with 25 μM cisplatin for 4 hrs. Scale bar: 10 μm. (G) Example STORM render image of γH2AX, 2 hrs after electroporation with Cas9 and 50A guide (GC-selective) (scale bar 2 μm). (H) Cluster map from panel F depicting density of γH2AX molecules per μm2. Clusters are defined by detecting a minimum of 3 molecules within a search area corresponding to the STORM localisation precision. The search area then propagates and a group of molecules is considered to be a cluster if at least 10 molecules are found.

The confocal analysis reinforces the ability of our method to induce a quantitatively variable level of DNA damage, depending on the chosen guide sequence. To further address this point, we performed single molecule localisation microscopy (STORM) analysis with the 50A guide 2-hour measurement (Figure 4G). This approach is equivalent to the widefield imaging performed in Figure 2 and 3, but with approximately five-fold increased resolution and the ability to count single molecules and quantify clusters ^18,19^. We detected γH2AX foci and then quantified the foci using cluster analysis (Figure 4H). Dense clusters were identified, as expected for γH2AX foci following DNA damage. We observed an average of 37 clusters cell^−1^, again significantly higher than standard widefield imaging and close to the predicted number of cut sites (Figure 4E).

Overall, the above results show that widefield imaging leads to merging of nearby cut sites and thus underestimation of the true number of foci. However, when comparing the number of foci detected in widefield imaging across all the different guides tested, we can observe a good linear correlation between the predicted cut number and the mean detected foci at all time points up to 2hrs (Figure 5). This is with the exception of 100B which did not vary across the time course. This correlation is critical to the dosage response given by using Cas9. Overall, as previously found in DiVA cells ^10^, the 2-hour time point appears optimal for obtaining the desired levels of DNA damage, representing the best trade-off between the kinetics of damage induction and repair.

**Fig.5.**
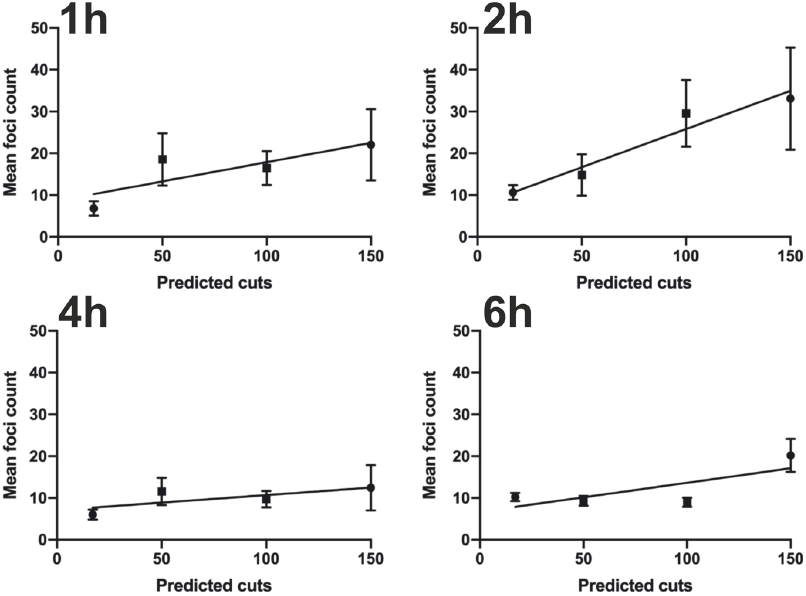
Correlation between the predicted number of cuts and detected γH2AX foci. Linear correlation plots of mean foci count per cell following electroporation with guide RNA to induce 17, 50, 100 and 150 cuts. The time refers to hours post-electroporation. The bars represent the 95% CI.

### Tuning the amount of DNA damage using Cas9 and promiscuous guides

#### (II) Intensity-based quantification of DNA damage

Particle analysis has optical limitations relating to the detection of individual foci. As a second independent approach to damage quantitation, we investigated the mean γH2AX fluorescence intensity across nuclei for each of the selected guide RNAs. This removes the need for resolving individual DSBs and instead provides an overall aggregate measure of the level of DNA damage and γH2AX signal.

As expected, mean intensity (Figure 6A) was significantly higher than the control measurements for all guide RNAs tested (Figure 1E). Moreover, there was once again a temporal increase in mean intensity suggesting damage accumulates up to 4 hours and begins to decrease, depending on the guide. Interestingly, the centromere guide can be distinguished from the HS17 guide at 2 hours and it appears that the damage is largely repaired by 6 hours. This is consistent with a resolution limitation when performing the single particle analysis based on foci counting. Consistent with the prior data, using guide RNAs with increasing predicted cuts led to an increase in total γH2AX staining intensity. However, exploring the data more closely shows that the 150 guide is an outlier because the intensity cannot be distinguished from the other guides. This may suggest that the nuclease activity is not efficient with the 150 guide RNA.

**Fig.6.**
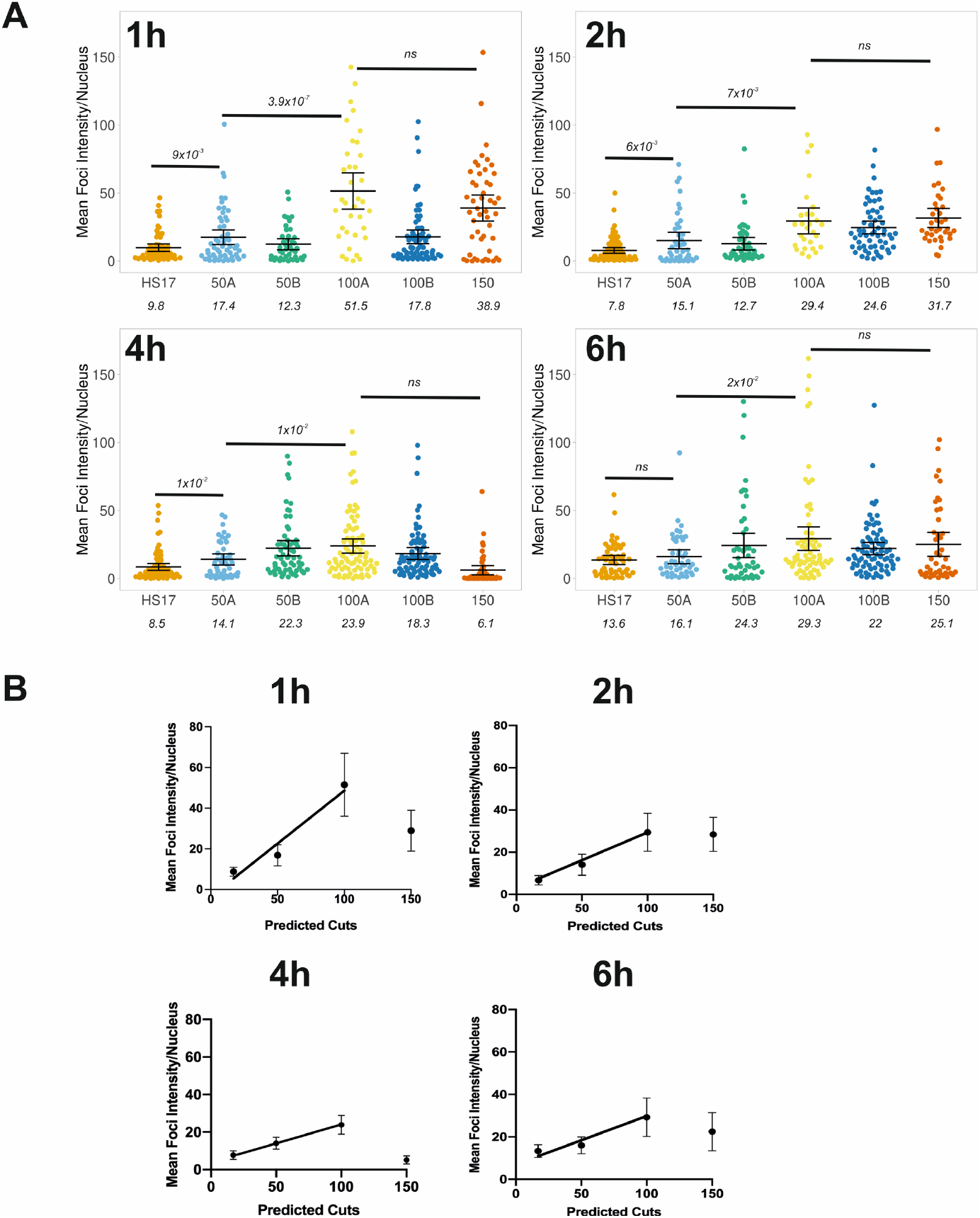
Correlation between the predicted number of cuts and γH2AX fluorescence intensity. (A) Quantitative image analysis, as described in the methods, from the experiments in Fig.2. The Mean intensity per nucleus is plotted for each guide RNA. Each data point is an individual nucleus. The p value is calculated from a two-tailed t-test and the error bars represent sd. The mean values are presented. The time refers to hours post-electroporation. (B) Linear correlation plots of mean intensity per nuclei following electroporation with guide RNA to induce 17, 50, 100 and 150 cuts. The bars represent the 95% CI. The data for 150 did not vary across the time course so it was removed from the correlation fitting.

The number of foci subsided quickly after the 2-hour measurements, yet the intensity measurements do not follow this trend. It is therefore possible that foci are fusing for repair, thus reducing the overall number of foci but maintaining overall γH2AX levels. This once again highlights the need for caution when only applying particle analysis.

There was a correlation between expected cuts and mean intensity for up to 100 cuts, as with the particle detection (Figure 6B). This reinforces the lack of effective cutting with the 150 guide because the intensity was lower than expected in all time points. Nevertheless, the positive correlation, up to 100 cuts, was observed across all time points which highlights the gain of using intensity over particle detection methods. It also implies there are foci fusion events occurring as repair occurs during the latter time points.

## DISCUSSION

We have described that by using a programmable Cas9 system it is possible to induce and detect targeted and titratable DNA damage (DSBs) and quantitate this via γH2AX immunostaining. This approach is an expansion upon the work of van der Berg *et al.* ^16^ in an attempt to establish a functional proof of concept for designing crRNAs that have multiple recognition sites for DSB-induction at lower logistical cost.

The electroporation of large ribo-protein complexes such as Cas9-guide RNA has high versatility and can be applied to almost any cell line without any prior cell line engineering. Importantly, electroporation does not cause stress or damage resulting in false-positives. We acknowledge that engineered cell lines which stably express Cas9 and the guides will have a greater damage induction efficiency within a cell population. However, generating engineered cell lines at scale is not practical, and thus this approach cannot be used for (e.g.) high-throughput comparison of DNA damage responses across different cell types. In contrast, our approach allows targeted DNA damage induction in any electroporatable cell line and will enable a wider understanding of how DNA damage repair varies in different genomic contexts, different cell types, and with differing numbers of breaks per cell.

We implemented two image analysis-based approaches to determine the efficiency of damage induction. As expected, due to the additive nature of the fluorescence signal, intensity measurements were more robust to determine differences between the guide RNAs. Particle detection, with the aim of directly counting damage sites, was dependent upon the resolution of the microscopy technique employed.

Our data showed that the γH2AX response is linear up to 100 cuts but it is likely that not all potential sites are cut by Cas9. By using a cocktail of several crRNAs with fewer cuts each, similarly to Zhou *et al*, it might be possible to specify even more precisely the number of cuts induced in any given experiment 20. However, this comes with a trade-off in that increasing the number of different guides used will decrease the concentration of any given guide delivered to each cell. This may reduce cutting efficiency, as well as increasing the individual cell variability depending on the precise amounts of the different guides taken up by each cell.

Overall, therefore, our approach of using promiscuous guide RNAs represents an important new means to target DNA damage to a range of desired genomic regions in an inducible and titratable manner. This will add to the repertoire of techniques available to dissect the sophisticated DNA damage response across cell types and chromatin contexts.

## ACKNOWLEDGEMENTS

We thank the UKRI-MRC (MR/M020606/1) and UKRI-STFC (19130001) to C.P.T. We also thank Darren Griffin and Anastasios Tsaousis (University of Kent) for sharing of equipment, and Dr Ben Skinner (University of Essex) and Dr Aaron McKenna (University of Washington) for assisting with the *in silico* design of the guide RNAs.

## AUTHOR CONTRIBUTIONS

C.P.T. and P.I.J.E conceived the study. I.E. C.P.T. and P.I.J.E designed experiments. I.E. performed experiments. I.E. and C.P.T. analysed data. Y.H-G. performed cell culture and assistance with electroporation. N.F. supported confocal microscopy measurements. A.W.C purified the dCas9 protein. A.dS. and A.W.C. performed experiments with cisplatin. Super Resolution Imaging was supported by L.W. and M.M-F. C.P.T. and P.I.J.E supervised the study. C.P.T. and P.I.J.E wrote the manuscript with comments from all authors.

## Competing financial interests

The authors declare no competing financial interests.

## MATERIALS AND METHODS

### Constructs

Plasmid pET-Cas9-NLS-6xHis responsible for encoding SpCas9 (wild type Cas9 derived from S. pyogenes) containing nuclear localisation signal (NLS) sequence and 6x-Histidine tag fused at the C-terminal, was obtained from Addgene (#62933). Plasmid pET302-6His-dCas9-Halo, the equivalent nuclease-deficient Cas9 was obtained from Addgene (#72269). All plasmids were verified by DNA sequencing.

### Protein expression and purification in Escherichia coli

Recombinant constructs were expressed in *E.coli* BL21 DE3 cells (Invitrogen) in Luria Bertani media. Proteins were purified by affinity chromatography (HisTrap FF, GE Healthcare). The purest fractions were then further purified through a Superdex 200 16/600 column (GE Healthcare).

### Cell culture

MCF10a (ATCC CRL-10317) cells were cultured at 37°C and 5% CO_2_, in 50% Gibco MEM Alpha medium with GlutaMAX (no nucleosides) and 50% Ham’s F-12 nutrient mixture, supplemented with 5% Fetal Bovine Serum (Gibco), 5% horse serum, penicillin-streptomycin mix diluted to 100 units mL-1, 50μg Cholera toxin, 5μg insulin, 20ngmL-1 human EGF and 0.5μgmL-1 hydrocortisone, 100 units/ml penicillin and 100 μg/ml streptomycin (Gibco).

### Drug Treatment

Cisplatin [*cis*-diammineplatinum(II) dichloride] (Sigma) was resuspended in a 0.9% NaCl solution to a concentration of 3.3 mM, following manufacturer’s instructions and used at a concentration of 25 μM.

### Electroporation

MCF10A cells were harvested with 0.25% trypsin/EDTA centrifugation at 500 rpm 4°C. The cells were then resuspended in 37°C Opti-MEM Reduced Serum Medium (Gibco).

Both crRNA (Dharmacon) and tracrRNA (Dharmacon Edit-R CRISPR-Cas9 Synthetic tracrRNA – U-002005-20) stock solutions (200μM each) prepared by adding the appropriate volume of RNase-free water. Then, the 100μM solution of crRNA:tracrRNA duplex was created by combining 200μM stock solutions in a 1:1 ratio. The solution was gently mixed for 10 min and stored at −20°C for future experiments. The project also utilised HS17 crRNA (5’-CAGACAGGCCCAGATTGAGG-3’) from Berg et al 16. The Cas9 ribonucleoprotein (RNP) complex was created by combining 1.5μM Cas9 protein and 3μM RNA final concentration and kept in ice until mixed with the resuspended cells in Opti-MEM medium.

Electroporation of the Cas9:RNA complex was achieved using a Gene Pulser/MicroPulser Electroporation Cuvettes with 0.2 cm gap cuvettes at in Gene Pulser Xcell Electroporation System and an exponential pulse at 300V and 300μF. Complete cell culture media was then added to the Opti-MEM in 1:1 ratio. Electroporated MCF10A cells were seeded on to coverslips pre-coated with 50 μg/ml poly-D-lysine (Sigma) and incubated at 37°C in 5% CO_2_.

### Immunofluorescence

MCF10a cells were fixed for 15 min at room temperature in 4% (w/v) paraformaldehyde (PFA) in TBS and residual PFA was quenched for 15 min with 50 mM ammonium chloride in TBS. For staining with Halo-TMR (Promega G8252), 10 nM ligand was added for 15 min and washed three times in warm cell culture media before fixation. All subsequent steps were performed at room temperature. Cells were permeabilised and simultaneously blocked for 15 min with 0.1 % (v/v) Triton X-100 and 2 % (w/v) BSA in TBS. Cells were then immuno-stained by 1 h incubation with the indicated primary and subsequently the appropriate fluorophore-conjugated secondary antibody (details below), both diluted in 2 % (w/v) BSA in TBS. The following antibodies were used at the indicated dilutions: Rabbit anti-Cas9 (1:200, Abcam ab204448), Mouse anti-phospho-H2A.X (1:500 Sigma 05-636), Donkey anti-rabbit Alexa Fluor 488-conjugated (1:250, Abcam Ab181346), Donkey anti-mouse Alexa Fluor 647-conjugated (1:250, Abcam Ab150103). For Hoechst staining, coverslips were washed three times in TBS followed by Hoechst 33342 solution for 10 mins at RT in the dark. The coverslips were then washed three times in TBS and once with ddH_2_O. Coverslips were mounted on microscope slides with Mowiol (10% (w/v) Mowiol 4-88, 25% (w/v) glycerol, 0.2 M Tris-HCl, pH 8.5), supplemented with 2.5% (w/v) of the anti-fading reagent DABCO (Sigma).

### Widefield Fluorescent Imaging

Widefield immunofluorescence images were obtained using CytoVision Olympus BX61 microscope equipped with Olympus UPlanFL 100 X/1.30 NA oil objective lens and Hamamatsu Photonics Digital CCD Camera ORCA-R2 C10600-10B-H.

### Confocal Imaging

Cells were visualised using the ZEISS LSM 880 confocal microscope. This was equipped with a Plan-Apochromat 63x/1.4 NA oil immersion lens (Carl Zeiss, 420782-9900-000). The built-in dichroic mirrors (Carl Zeiss, MBS-405, MBS-488 and MBS-561) were used to reflect the excitation laser beams on to cell samples. The emission spectral bands for fluorescence collection were 410 nm-524 nm (Hoechst, Thermo Fisher), 493 nm-578 nm (AlexaFluor 488, Thermo Fisher) and 650 nm-697 nm (AlexaFluor 647, Thermo Fisher). The detectors consisted of two multi anode photomultiplier tubes (MA-PMT) and 1 gallium arsenide phosphide (GaAsP) detector. The green channel was imaged using GaAsP detector, while the blue and red channels were imaged using MA-PMTs. ZEN software (Carl Zeiss, ZEN 2.3) was used to acquire and render the confocal images.

### Image Analysis

For single particle detection, we used Fiji 21 to split the RGB channels and convert the γH2AX channel to a binary image. The Despeckle function was used to remove background noise from the images. The area of the nucleus was selected by creating a mask from the Hoechst channel. The Analyze Particles function used to calculate the number of foci in a given nucleus. For fluorescence intensity measurements. The binary image for single particle detection was used to create a mask and then the mean pixel intensity was calculated for each particle.

### STORM Imaging

Cells were seeded on pre-cleaned No. 1.5, 25-mm round glass coverslips, placed in 6-well cell culture dishes. Glass coverslips were cleaned by incubating them for 3 hours, in etch solution, made of 5:1:1 ratio of H_2_O : H_2_O_2_ (50 wt. % in H_2_O, stabilized, Fisher Scientific) : NH_4_OH (ACS reagent, 28-30% NH_3_ basis, Sigma), placed in a 70°C water bath. Cleaned coverslips were repeatedly washed in filtered water and then ethanol, dried and used for cell seeding. Cells were fixed in pre-warmed 4% (w/v) Paraformaldehyde (PFA) in PBS and residual PFA was quenched for 15 min with 50 mM ammonium chloride in PBS. Immunofluorescence was performed in filtered sterilised TBS. Cells were permeabilized and simultaneously blocked for 30 min with 3% (w/v) BSA in TBS, supplemented with 0.1 % (v/v) Triton X-100. Permeabilized cells were incubated for 1h with the primary antibody and subsequently the appropriate fluorophore-conjugated secondary antibody, at the desired dilution in 3% (w/v) BSA, 0.1% (v/v) Triton X-100 in TBS. The antibody dilutions used were the same as for the normal IF protocol (see above), except from the secondary antibodies which were used at 1:250 dilution. Following incubation with both primary and secondary antibodies, cells were washed 3 times, for 10 min per wash, with 0.2% (w/v) BSA, 0.05% (v/v) Triton X-100 in TBS. Cells were further washed in PBS and fixed for a second time with pre-warmed 4% (w/v) PFA in PBS for 10 min. Cells were washed in PBS and stored at 4 °C, in the dark, in 0.02% NaN3 in PBS, before proceeding to STORM imaging.

Before imaging, coverslips were assembled into the Attofluor^®^ cell chambers (Invitrogen). Imaging was performed in freshly made STORM buffer consisting of 10 % (w/v) glucose, 10 mM NaCl, 50 mM Tris - pH 8.0, supplemented with 0.1 % (v/v) 2-mercaptoethanol and 0.1 % (v/v) pre-made GLOX solution which was stored at 4 0C for up to a week (5.6 % (w/v) glucose oxidase and 3.4 mg/ml catalase in 50 mM NaCl, 10 mM Tris - pH 8.0). All chemicals were purchased from Sigma. Imaging was undertaken using the Zeiss Elyra PS.1 system. Illumination was from a HR Diode 642 nm (100 mW) lasers where power density on the sample was 7-12 kW/cm^2^.

Imaging was performed under highly inclined and laminated optical (HILO) illumination to reduce the background fluorescence with a 100x/ 1.46NA oil immersion objective lens (Zeiss alpha Plan-Apochromat) with a BP 420-480/BP495-550/LP 650 filter. The final image was projected on an Andor iXon EMCCD camera with 25 msec exposure for 20000 frames. The focal plane was locked using Definite Focus function in the microscope during image acquisition.

The images were processed through our STORM analysis pipeline using the Zeiss Zen Black software. Single molecule detection and localisation was performed using a 9-pixel mask with a signal to noise ratio of 6 in the “Peak finder” settings while applying the “Account for overlap” function. This function allows multi-object fitting to localise molecules within a dense environment. Molecules were then localised by fitting to a 2D Gaussian.

The render was then subjected to model-based cross-correlation lateral drift correction and detection grouping to remove detections within multiple frames. Typical localisation precision was 20 nm for Alexa-Fluor 647. The final render was then generated at 10 nm/pixel and displayed in Gauss mode where each localisation is presented as a 2D gaussian with a standard deviation based on its precision. The localisation table was exported as a csv for import in to Clus-DoC.

### Clus-DoC

The single molecule positions were exported from Zeiss Zen Black and imported into the Clus-DoC analysis software ^22^ (https://github.com/PRNicovich/ClusDoC). The region of interest was determined by the nuclear staining. First the Ripley K function was completed to identify the r max. The r max was then assigned for DBSCAN. The MinPts was 3 and a cluster required 10 locations, with smoothing set at 7 nm and epsilon set at the mean localization precision for the dye. All other analyses parameters remained at default settings 22. Data concerning each cluster was exported and graphed using Plots of Data 23.

### In silico design of guide RNAs

The crRNAs used in this project were designed using FlashFry developed by McKenna, A. and Shendure, J. ^17^. FlashFry was downloaded from Github - https://github.com/mckennalab/FlashFry and configured according to the author’s recommendations. The binary database was created based on the latest human genome (hg38 build) in FASTA format from UCSC. The verification of the newly designed crRNA hits across the human genome was done in BLAST/BLAT search from Ensembl with adjusted option to report the maximum number of hits to report to 5000, E-value for alignment report at 1.0, match/mismatch scores equal to 1,-1 with filtering low complexity regions and query sequences options enabled.

### Graphics

Unless stated, data fitting and plotting was performed using Plots of data 23 and GraphPad. Cartoons were generated using the BioRender software.

## Data Availability

The data supporting the findings of this study are available from the corresponding author on request.

## Notes

### Competing Interest Statement

The authors have declared no competing interest.

