## Supplementary Figures for "A targeted and tuneable DNA damage tool using CRISPR/Cas9"

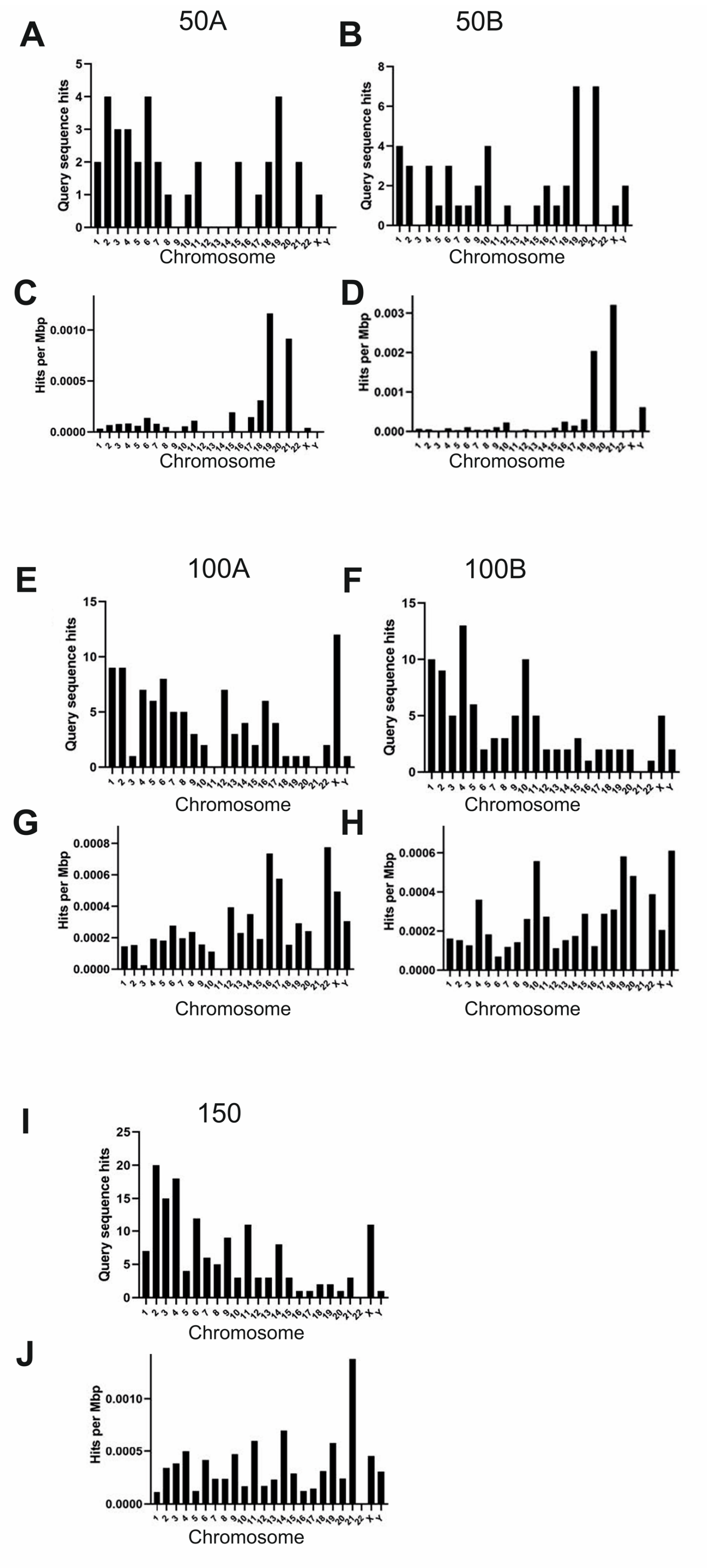


**Supplementary Figure 1. Overview of genomic targeting for the crRNA sequences.** Data are presented as Number of sequence hits (potential cuts) across the chromosomes and Hits per Mbp across the chromosomes for each of the crRNA guides designed in this study.
